## Supplemental information for "Convergent evolution of distinct D-ribulose utilisation pathways in attaching and effacing pathogens"

**Supplementary Figures 1-10**

**Supplementary table 1** – Bacterial strains used in this study.

**Supplementary table 2** – Primers used in this study.

**Supplementary table 3** – Plasmids used in this study.

**Supplementary table 4 –** X-ray data collection and refinement statistics

**Supplementary references**

**Uncropped images**

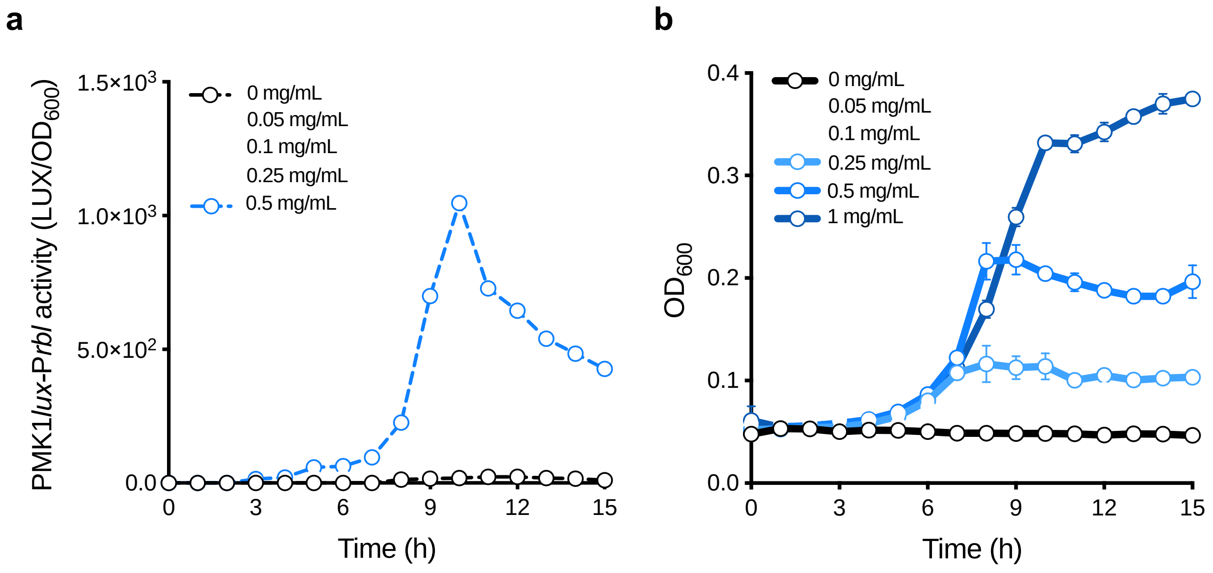

**Supplementary Fig. 1. The *C. rodentium ROD_24811-61* locus is responsive to a range of D-ribulose concentrations. a,** Growth analysis of wild type *C. rodentium* in M9 minimal media supplemented with the indicated concentration range of D-ribulose as a sole carbon source. **b,** Transcriptional reporter assay of *C. rodentium* transformed with the pMK1*lux*-P$_{24811}$ plasmid cultured in MEM-HEPES alone or supplemented with the indicated concentration range D-ribulose. Error bars for reporter assays and growth curves represent the standard deviation from three independent experiments (*n* = 3 biological replicates).

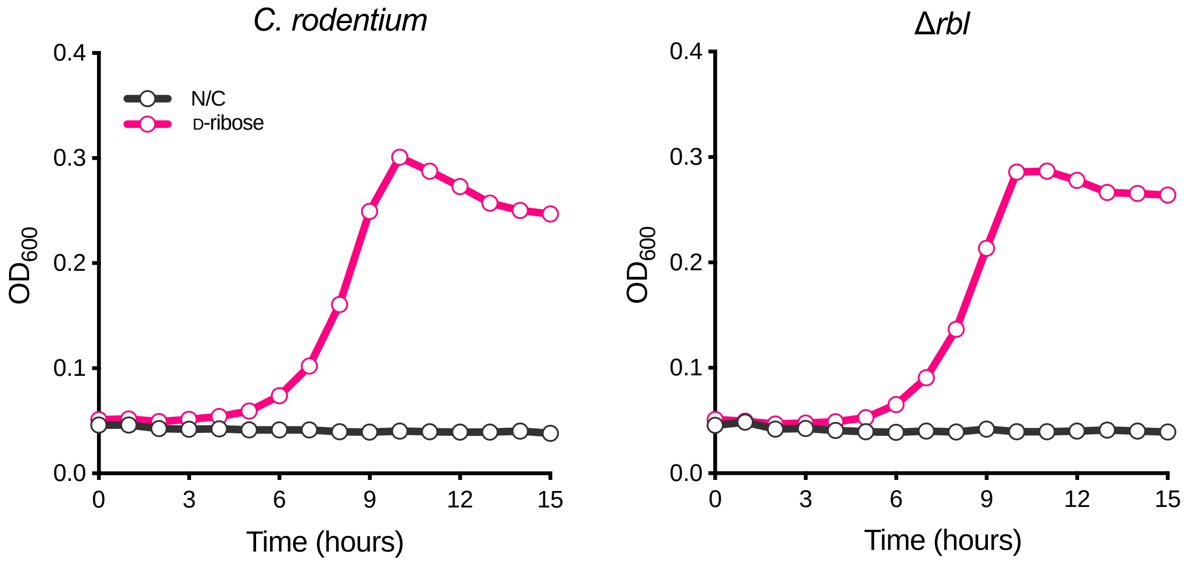

**Supplementary Fig. 2. The *C. rodentium ROD_24811-61* locus is not required for growth on D-ribose**. Growth analysis of wild type *C. rodentium* and Δ*rbl* cultured in M9 minimal media supplemented with 0.5 mg/mL D-ribose. The no sugar control indicates wild type *C. rodentium* inoculated into M9 without a carbon source. Error bars represent the standard deviation from three independent experiments ($n$ = 3 biological replicates).

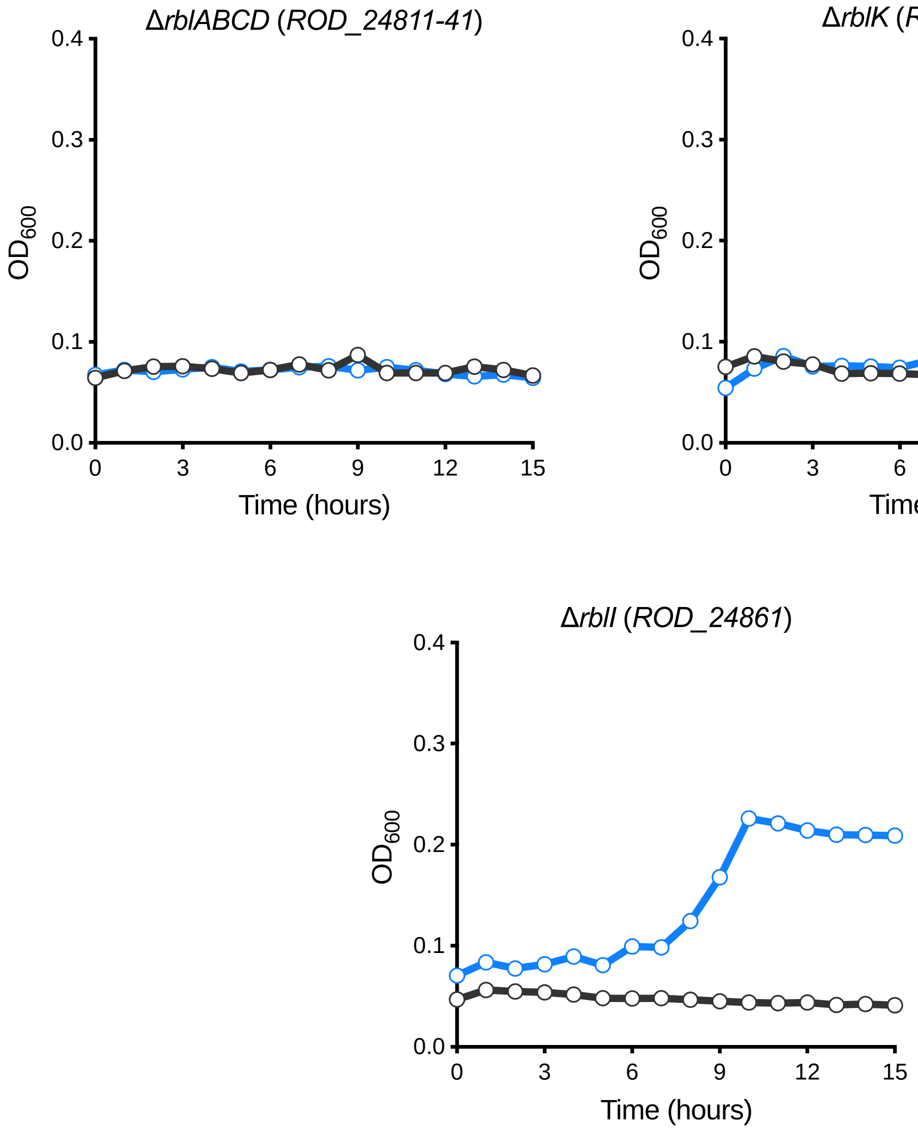

**Supplementary Fig. 3. The Rbl ABC transporter and D-ribulokinase are required for utilisation of D-ribulose by *C. rodentium*.** Growth analysis of wild type *C. rodentium, ΔrblABCD, ΔrblK* and *ΔrblI* cultured in M9 minimal media supplemented with 0.5 mg/mL D-ribose. The no sugar control indicates wild type *C. rodentium* inoculated into M9 without a carbon source. Error bars represent the standard deviation from three independent experiments (*n* = 3 biological replicates).

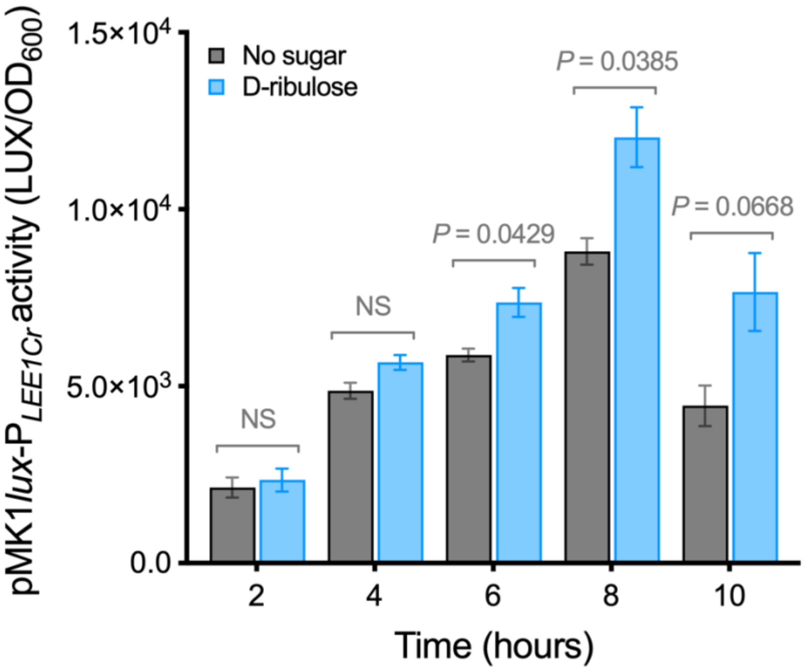

**Supplementary Fig. 4. D-ribulose metabolism enhances expression of the LEE-encoded type 3 secretion system in *C. rodentium*.** Transcriptional reporter analysis of *C. rodentium* transformed with pMK1*lux*-$P_{LEE1Cr}$ cultured in MEM-HEPES (grey) or supplemented with 0.5 mg/mL of D-ribulose (blue). Data are depicted as luminescence units (LUX) divided by optical density ($OD_{600}$) at each timepoint. Statistical significance was determined by two-tailed students' *t*-test. Error bars represent standard deviation (*n* = 2 biological replicates).

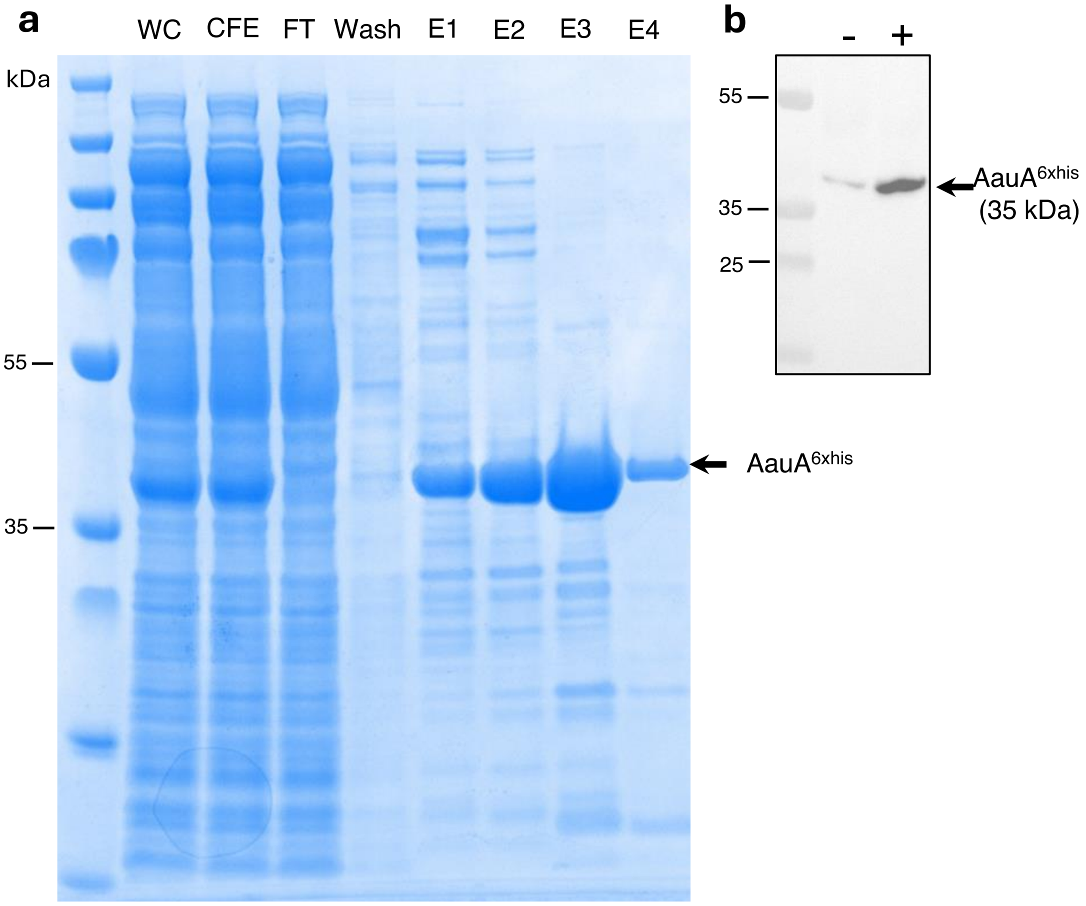

**Supplementary Fig. 5. Overexpression and purification of recombinant AauA from EHEC. a**, SDS-PAGE analysis of AauA with C-terminal His-tag purification by immobilized metal affinity chromatography. WC: Whole cell fraction; CFE: Cell free extract; FT: Flow through. E1 and E2 correspond to elution with 10 mM imidazole. E3 and E4 correspond to elution with 100 mM imidazole. **b,** Western blot detection of his-tagged AauA before and after induction with 0.5 mM IPTG.

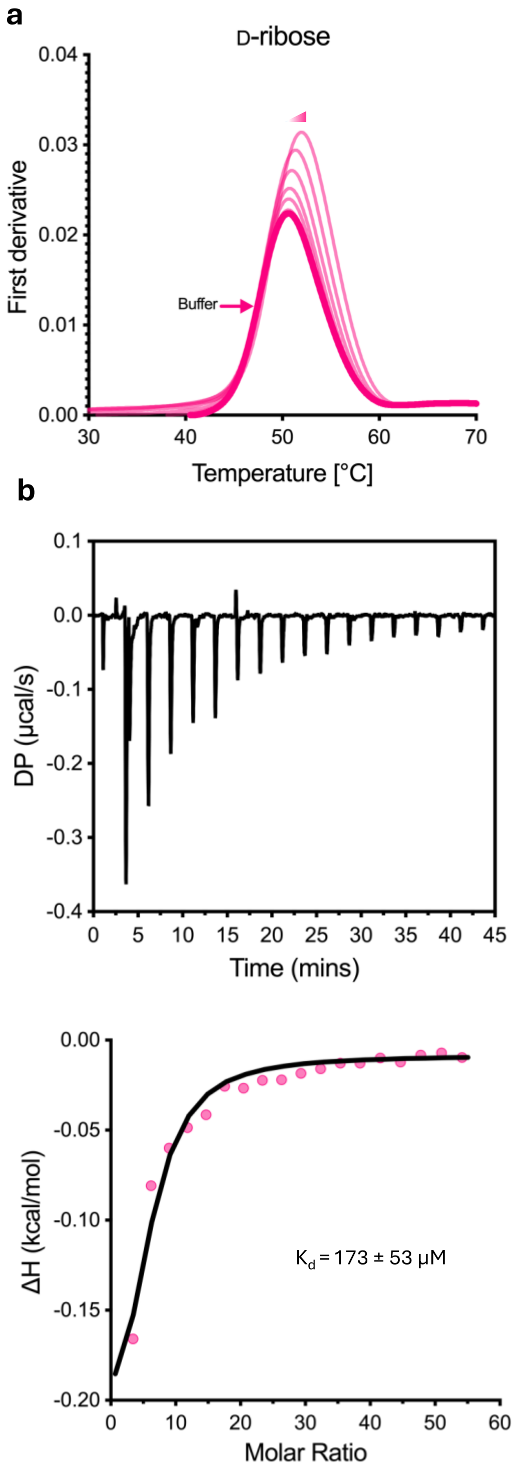

**Supplementary Fig. 6. AauA shows low affinity for D-ribose. a,** NanoDSF data depicting the shift in melting temperature ($\Delta T_m$) of purified AauA in the presence of increasing concentrations of D-ribose (pink). The buffer only control is illustrated in bold. NanoDSF melting experiments were performed in technical triplicate ($n$ = 3). **b,** Representative ITC thermogram of D-ribose titrated into purified AauA. The integration of heats derivative curve is illustrated on the bottom panel, with the corresponding calculated $K_d$ shown.

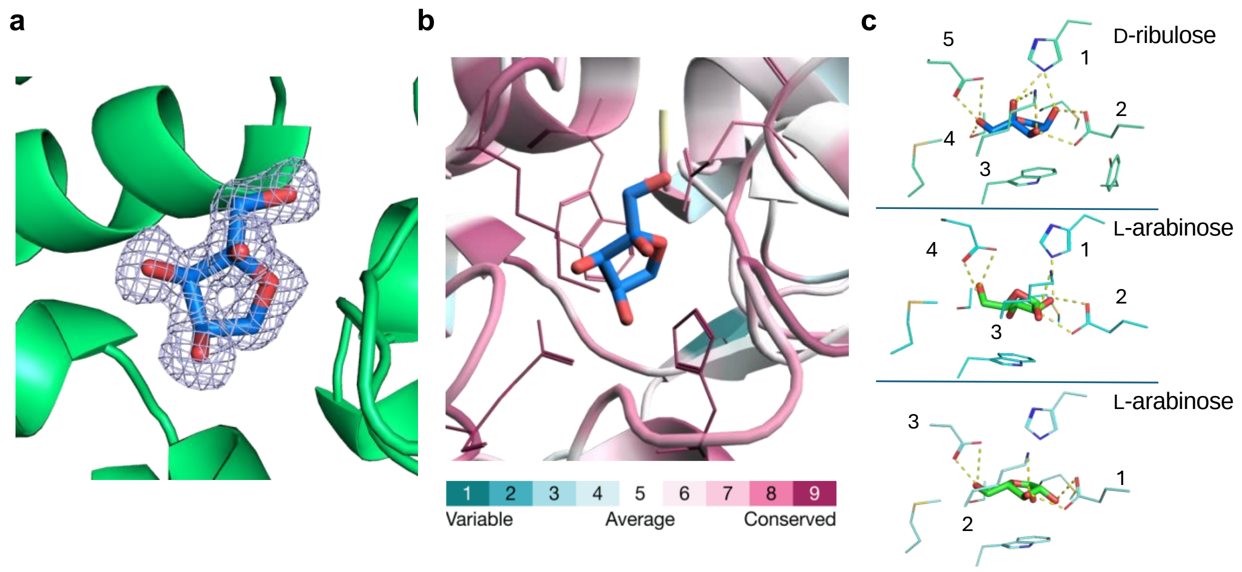

**Supplementary Fig. 7. Conserved binding site of AauA coordinates D-ribulose with high specificity. a,** Polder omit map (contoured at 3.5 σ) showing the conformation of alpha-D-ribulose within the binding pocket of AauA. **b,** Highly conserved nature of all coordinating residues within the substrate-binding site as analysed using ConSurf. **c,** High complementarity of the AauA binding towards alpha-D-ribulose over L-arabinose. AauA has 5 residues that can hydrogen bond with alpha-D-ribulose (top structure) versus 4 residues and 2 putative L-arabinose conformations (middle and bottom panels).

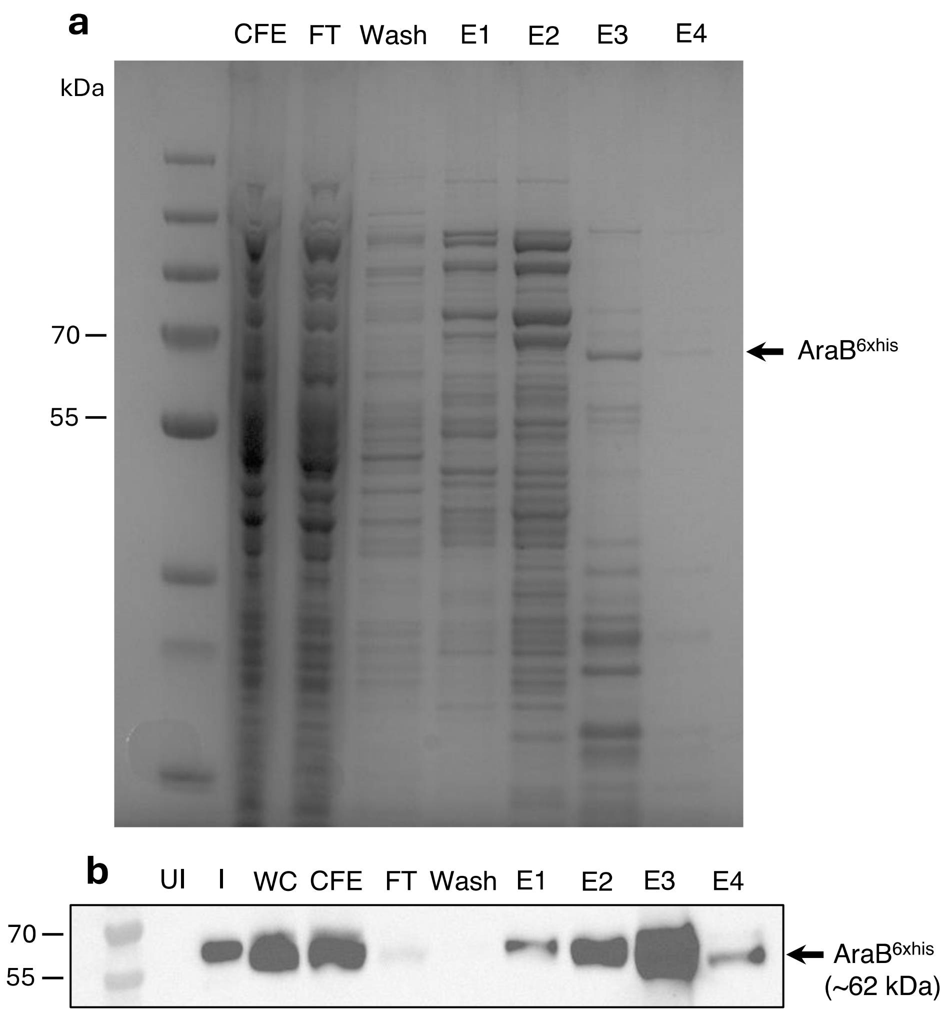

**Supplementary Fig. 8. Overexpression and purification of recombinant AraB from EHEC. a**, SDS-PAGE analysis of AraB with C-terminal His-tag purification by immobilized metal affinity chromatography. WC: Whole cell fraction; CFE: Cell free extract; FT: Flow through. E1 and E2 correspond to elution with 10 mM imidazole. E3 and E4 correspond to elution with 100 mM imidazole. **b,** Western blot detection of his-tagged AraB from the same fractions in panel **a**. Uninduced (UI) and induced (I) fractions are also shown.

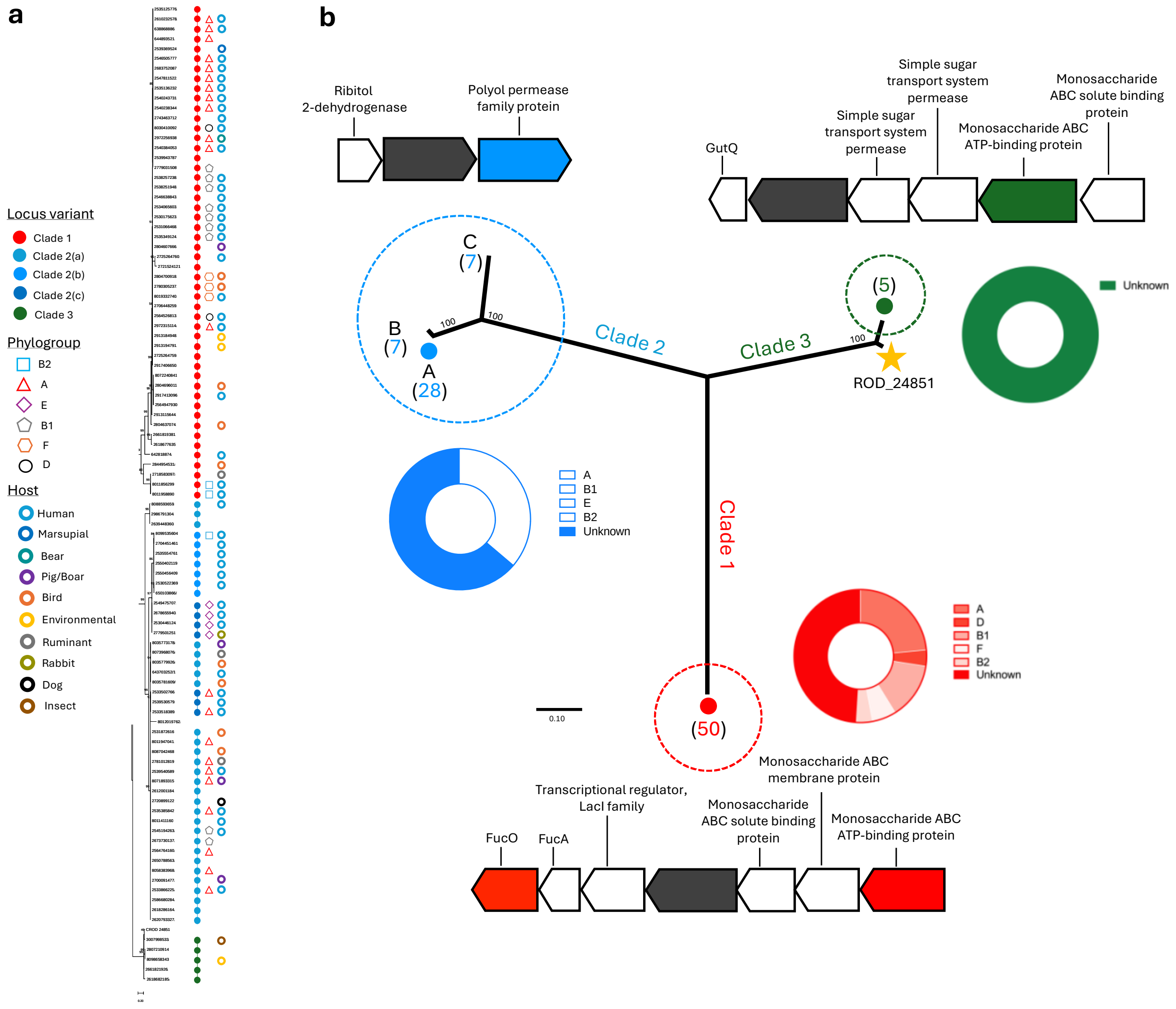

**Supplementary Fig. 9. Analysis of D-ribulokinase carriage in *Escherichia* spp. a**, Maximum likelihood phylogenetic tree of identified FGGY-family carbohydrate kinase (TIGR01315) homologues across *E. coli*, representative of 100 bootstrap replicates. Associated strain metadata from the Integrated Microbial Genomes and Microbiomes and Enterobase databases is displayed. **b,** Maximum likelihood phylogeny highlighting the three clades identified amongst strains carrying a predicted D-ribulokinase. Representative strains from each clade were used to generate the phylogeny. The organisation of loci associated with D-ribulokinase is shown, and the phylogroups strains were found to belong are summarised across each clade.

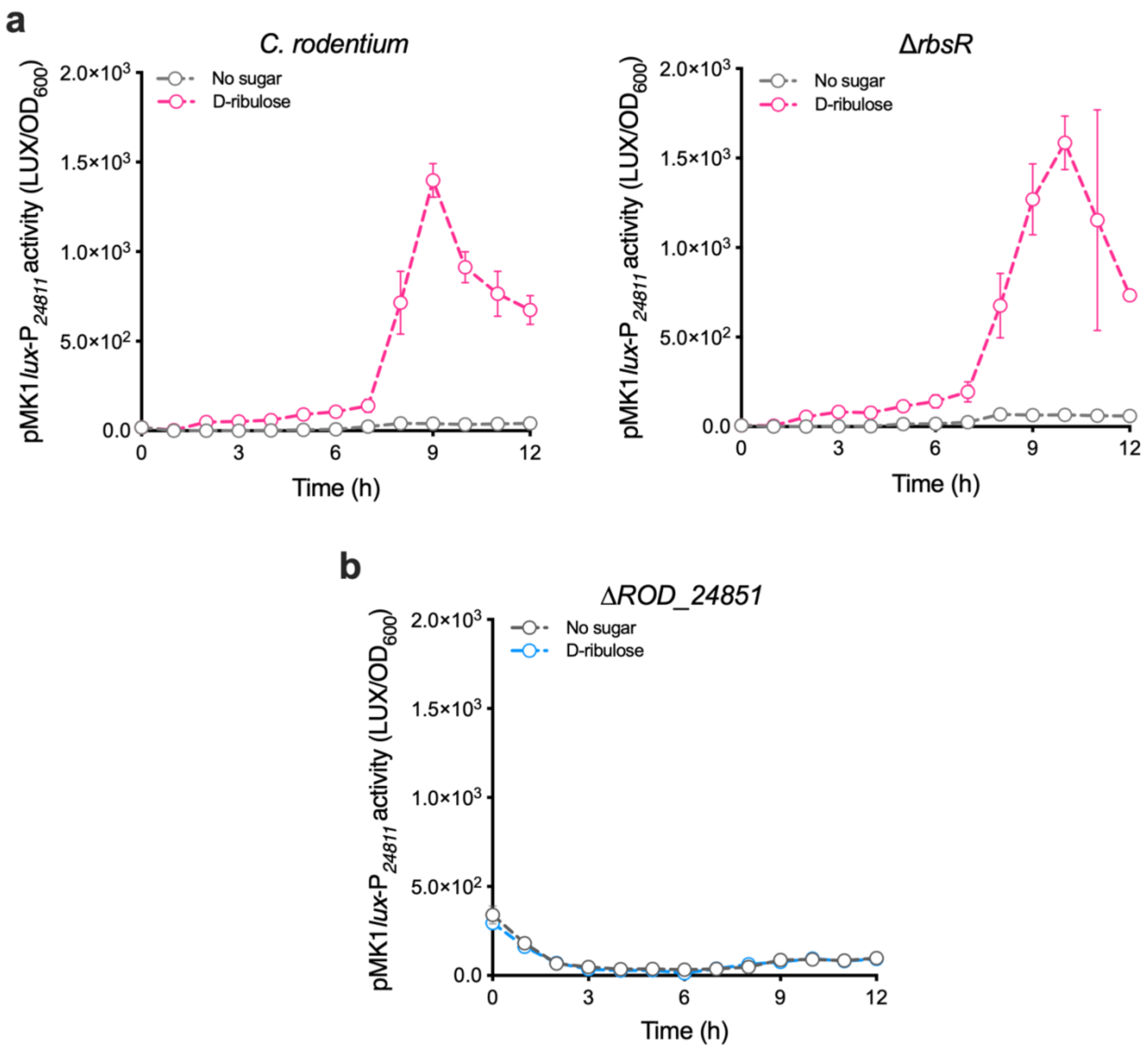

**Supplementary Fig. 10. D-ribulose metabolism is required to activate transcription of *rbl*. a,** Transcriptional reporter assay of wild type *C. rodentium* (left) and Δ*rbsR* (right) transformed with the pMK1*lux*-$P_{24811}$ plasmid cultured in MEM-HEPES alone (grey) or supplemented with 0.5 mg/mL D-ribulose. **b,** pMK1*lux*-$P_{24811}$ reporter assay in the *C. rodentium* Δ*rblK* mutant background, cultured in MEM-HEPES alone (grey) or supplemented with 0.5 mg/mL D-ribulose. Error bars for reporter assays represent the standard deviation from three independent experiments (*n* = 3 biological replicates).

**Supplementary Table 1** – Bacterial strains used in this study.

| Strain | Description | Reference |
|---|---|---|
| ZAP193 | EHEC O157:H7 str. ZAP193 (NCTC 12900) $Stx^-$ | Roe *et al*. 2004 |
| Δ*aau* | ZAP193 *Z0415-9* deletion mutant; $Kan^R$ $Strep^R$ | Cottam *et al*. 2024 |
| TUV93-0 | EHEC O157:H7 str. EDL933 $Stx^-$ | Campellone *et al*. 2002 |
| Δ*araBAD* | TUV93-0 *araBAD* deletion mutant; $Kan^R$ | Cottam *et al*. 2024 |
| ICC169 | *C. rodentium* O152 serotype; $Nal^R$ | Petty *et al*. 2010 |
| Δ*rbl* | ICC169 *rblABCDKI* (*ROD_24811-61*) deletion mutant; $Kan^R$ | This study |
| Δ*rblABCD* | ICC169 *ROD_24811-41* deletion mutant; $Kan^R$ | This study |
| Δ*rblK* | ICC169 *ROD_24851* deletion mutant; $Cm^R$ | This study |
| Δ*rblI* | ICC169 *ROD_24861* deletion mutant; $Kan^R$ | This study |
| Δ*rbsR* | ICC168 *rbsR* deletion mutant; $Kan^R$ | This study |

**Supplementary Table 2** – Primers used in this study.

| Primer Name | Description | Sequence |
|---|---|---|
| *ROD24811-51*_LRed_Fwd | ROD24811-51 KO forward primer | ttgctaacctcgtttcgtgacatgccctggtcccattaaaaggaacgacagtgtaggctggagctgcttc |
| *ROD24811-51*_LRed_Rev | ROD24811-51 KO reverse primer | cgcttcgctatagagccgccaggtggcggtggcctgctgccatgcgctgccatatgaatatcctccttag |
| *ROD24811-51*_Check_Fwd | ROD24811-51 KO check forward primer | tgccctggtcccattaaa |
| *ROD24811-51*_Check_Rev | ROD24811-51 KO check reverse primer | attaacgcctgccactgc |
| *ROD24811-41*_LRed_Fwd | ROD24811-41 KO forward primer | ttgctaacctcgtttcgtgacatgccctggtcccattaaaaggaacgacagtgtaggctggagctgcttc |
| *ROD24811-41*_LRed_Rev | ROD24811-41 KO reverse primer | ttcctacatcgacaccaataaaataactcgccatcatttttctcccgaaacatatgaatatcctccttag |
| *ROD24811-41*_Check_Fwd | ROD24811-41 KO check forward primer | acatgccctggtcccattaa |
| *ROD24811-41*_Check_Rev | ROD24811-41 KO check reverse primer | ggtaaatttcaatggcgcgc |
| *ROD24851*_LRed_Fwd | ROD24851 KO forward primer | cctctttatcgcattacagaatcgtaaagcctgatttcgggagaaaaatggtgtaggctggagctgcttc |
| *ROD24851*_LRed_Rev | ROD24851 KO reverse primer | cgcttcgctatagagccgccaggtggcggtggcctgctgccatgcgctgccatatgaatatcctccttag |
| *ROD24851*_Check_Fwd | ROD24851 KO check forward primer | atacggcgagtcctatctgc |
| *ROD24851*_Check_Rev | ROD24851 KO check reverse primer | ttatcatcaggtcgtcggca |
| *ROD24861*_LRed_Fwd | ROD24861 KO forward primer | gagatgtatcaggatcacatgaagtaccgtcagctgatgcaggaggcgttgtgtaggctggagctgcttc |
| *ROD24861*_LRed_Rev | ROD24861 KO reverse primer | tattttctgcatatcggaaaaagccccgtctatgggacggggccaggccacatatgaatatcctccttag |
| *ROD24861*_Check_Fwd | ROD24861 KO check forward primer | cgcagaccaaccgcattaag |
| *ROD24861*_Check_Rev | ROD24861 KO check reverse primer | taacgtcaggattgcagggg |
| pMK1*lux*-PROD24811_Fwd | Forward primer for cloning ROD24811 promoter with EcoRI | cccgaattcctgccgcgactgctggca |
| pMK1*lux*-PROD24811_Rev | Reverse primer for cloning ROD24811 promoter with BamHI | cccggatccattgtcgttccttttaat |
| pMK1*lux*_Check_Fwd | Forward primer to check pMK1*lux* cloning | ctataaaaataggcgtatcac |
| pMK1*lux*_Check_Rev | Reverse primer to check pMK1*lux* cloning | ctggccgttaataatgaatg |
| pSU-PROM-*rbl*_Fwd | Rbl Gibson assembly forward primer | tctaccacagaggaggatccatgaaattcaaactcgcattactac |
| pSU-PROM-*rbl*_Rev | Rbl Gibson assembly reverse primer | ctcgaggggtcgactctagatcataacgcctcctgcatc |
| pSU-PROM_Check_Fwd | Forward primer to check pSU-PROM cloning | ctcttcgctattacgccagc |
| pSU-PROM_Check_Rev | Reverse primer to check pSU-PROM cloning | accctcatcagtgccaacat |
| pSU-PROM_Linear_Fwd | pSU-PROM linearisation forward primer | tctagactcgacccctcg |

| | | |
|---|---|---|
| pSU-PROM_Linear_Rev | pSU-PROM linearisation reverse primer | ggatcctcctctgtggtag |
| pET28a-araB_Fwd | Forward primer for cloning *araB* with X | ccatggatggcgattgcaattggcctcgattttggc |
| pET28a-araB_Rev | Reverse primer for cloning *araB* with X | ctcgagtagagtcggaacggcctgggcagcctgtgc |
| pET28a-aauA_Fwd | Forward primer for cloning *aauA* with X | ccatggatgatgaataaacgttttgttatc |
| pET28a-aauA_Rev | Reverse primer for cloning *aauA* with X | ctcgagataaagtgagtcgatattgtcttt |
| pET28a_Check_Fwd | Forward primer to check pET28a cloning | acccctcaagacccgtttag |
| pET28a_Check_Rev | Reverse primer to check pET28a cloning | atcggtgatgtcggcgatat |

**Supplementary Table 3** – Plasmids used in this study.

| Plasmid | Description | Reference |
|---|---|---|
| pMK1*lux* | pBR322 ori with the *luxCDABE* operon and MCS; $Amp^R$ | Karavolos *et al*. 2008 |
| pMK1*lux*-$P_{LEE1Cr}$ | pMK1*lux* with the ICC168 LEE1 promoter cloned into the MCS; $Amp^R$ | This study |
| pMK1*lux*-$P_{aau}$ | pMK1*lux* with the ZAP193 *aau* promoter cloned into MCS; $Amp^R$ | Cottam *et al*. 2024 |
| pMK1*lux*-$P_{24811}$ | pMK1*lux* with the ICC168 *rbl* promoter cloned into the MCS; $Amp^R$ | This study |
| pSUPROM | Cloning vector for expression under the TatA promoter; $Kan^R$ | Jack *et al*. 2004 |
| pSU-*rbl* | pSUPROM with the full *rbl* locus cloned into MCS; $Kan^R$ | This study |
| pSU-*aau* | pSUPROM with ZAP193 *aau* cloned into MCS; $Kan^R$ | Cottam *et al*. 2024 |
| pET28a | Expression vector for recombinant His-tagging; $Kan^R$ | Lab stock |
| pET28a-*aauA* | pET28a with *aauA* (ZAP193 locus tag 0432; excluding signal peptide) cloned into the MCS; $Kan^R$ | This study |
| pET28a-*araB* | pET28a with *araB* (from ZAP193) cloned into the MCS; $Kan^R$ | This study |
| pKD46 | Lambda Red recombinase expressing plasmid; temperature sensitive; $Amp^R$ | Datsenko and Wanner, 2000 |
| pKD3 | Template plasmid for Lambda Red mutagenesis; $Cm^R$ | Datsenko and Wanner, 2000 |
| pKD4 | Template plasmid for Lambda Red mutagenesis; $Kan^R$ | Datsenko and Wanner, 2000 |
| pCP20 | FLP recombinase expressing plasmid; temperature sensitive; $Amp^R$ | Datsenko and Wanner, 2000 |

**Supplementary Table 4** – X-ray data collection and refinement statistics.

| | **AauA dataset*<br>(PDBID: 9I1M)** |
|---|---|
| **Data collection:** | |
| Beamline | Diamond Light Source I03 |
| Space group | C2 |
| Cell dimensions | |
| *a*, *b*, *c* (Å) | 76.38, 66.94, 57.85 |
| α, β, γ (°) | 90, 94.45, 90 |
| Resolution (Å) | 38.9-1.35 (1.37-1.35) |
| $R_{pim}$ | 0.018 (0.098) |
| $R_{meas}$ | 0.033 (0.14) |
| $I/\sigma I$ | 29.8 (7.5) |
| $CC_{1/2}$ | 0.999 (0.98) |
| Completeness (%) | 79.1 (18.9) |
| Redundancy | 6.2 (3.2) |
| **Refinement:** | |
| Resolution (Å) | 1.35 |
| No. reflections | 50386 (574) |
| $R_{work}$ / $R_{free}$ | 0.126/0.143 |
| No. atoms | 5509 |
| Protein | 4464 |
| Ligand/ion | 74 |
| *B*-factors (Å$^2$) | |
| Protein | 12.7 |
| Ligand/ion | 23.26 |
| R.m.s. deviations | |
| Bond lengths (Å) | 0.0124 |
| Bond angles (°) | 2.022 |
| Rotamer outliers (%) | 0.42 |
| Ramachandran (%)<br>Favoured regions<br>Allowed regions<br>Outliers | <br>97.22<br>2.78<br>0 |
| Molprobity score | 1.66 |

*Values in parentheses are for highest-resolution shell.

**Uncropped images:**

Supplementary Figure 5a – Full SDS-PAGE gel

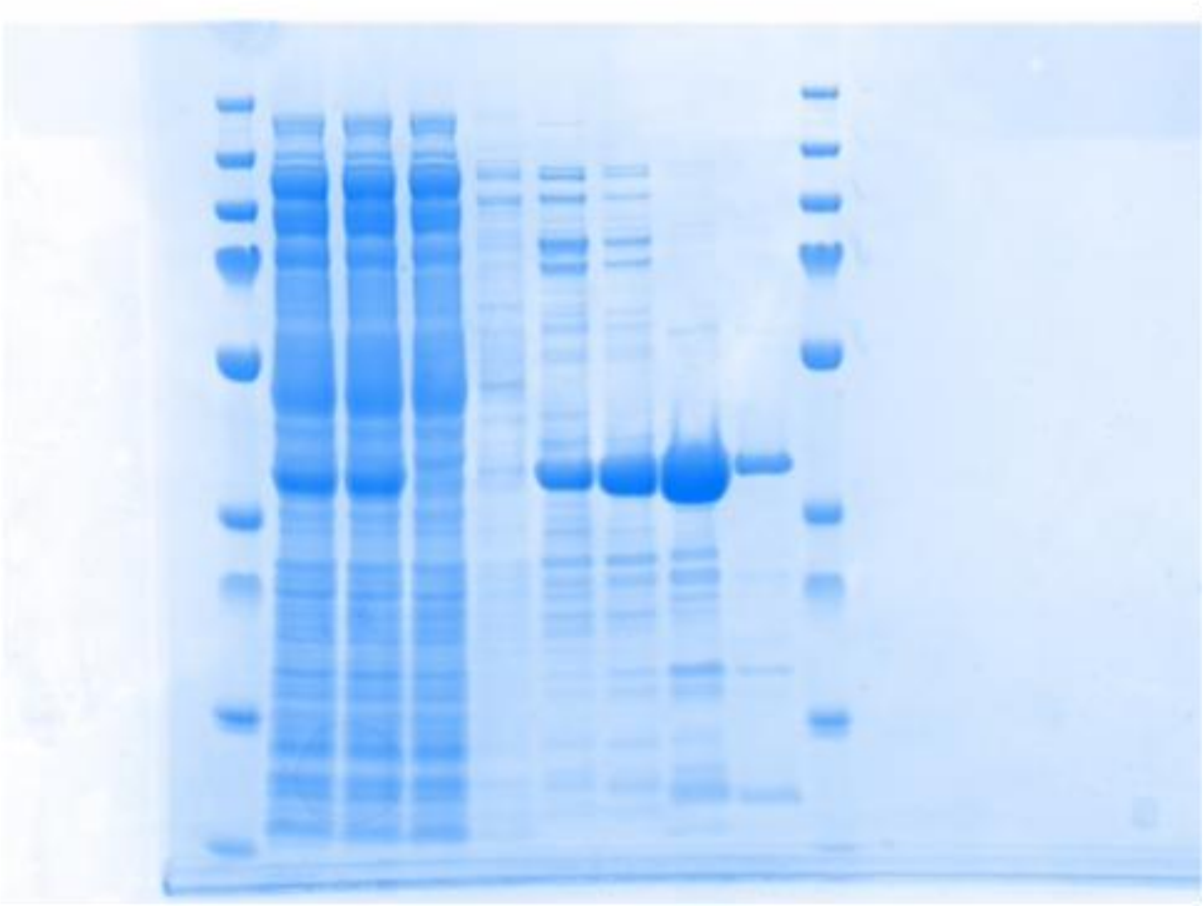

Supplementary Figure 5b – Full Western blot membrane

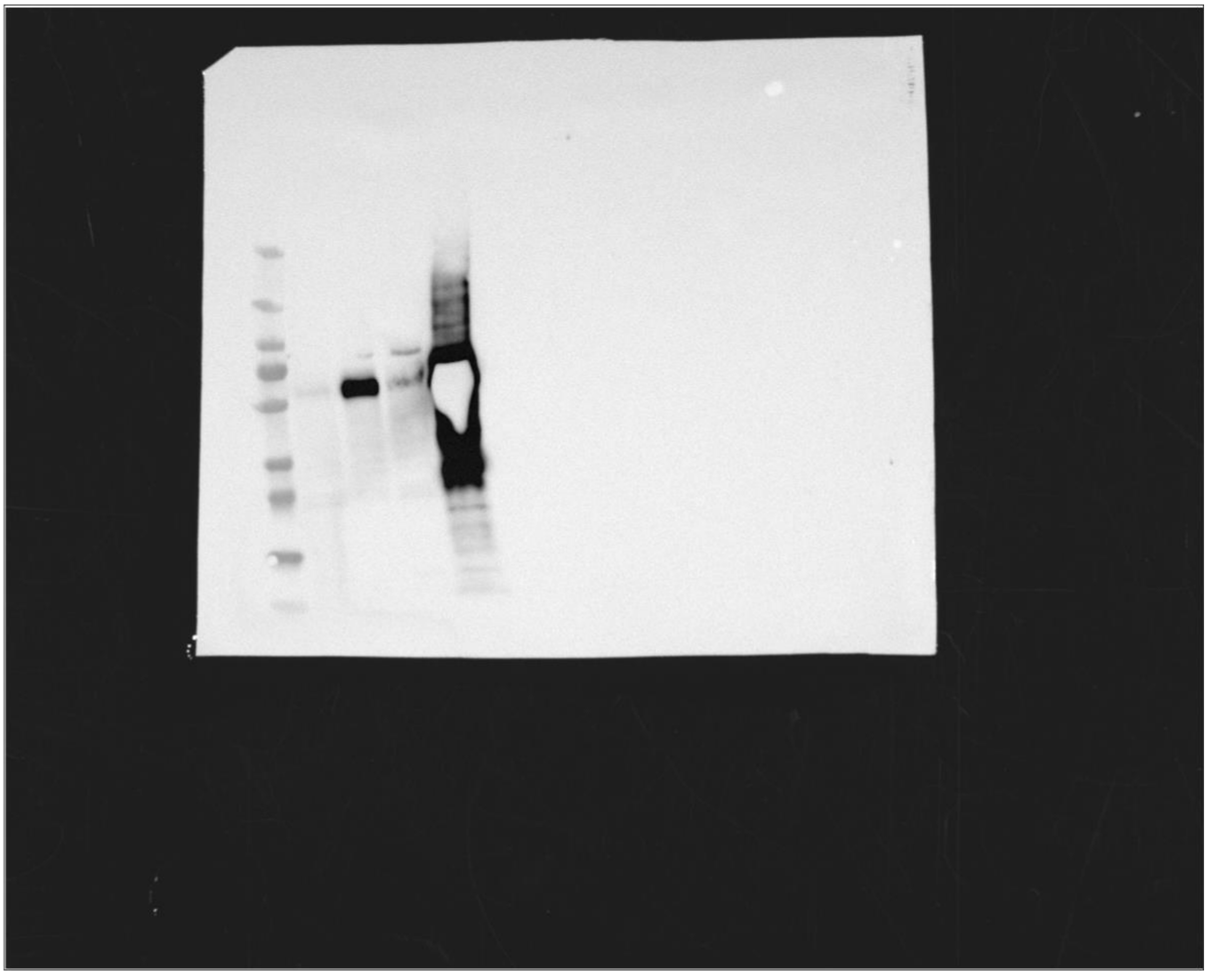

Supplementary Figure 8a – Full SDS-PAGE gel

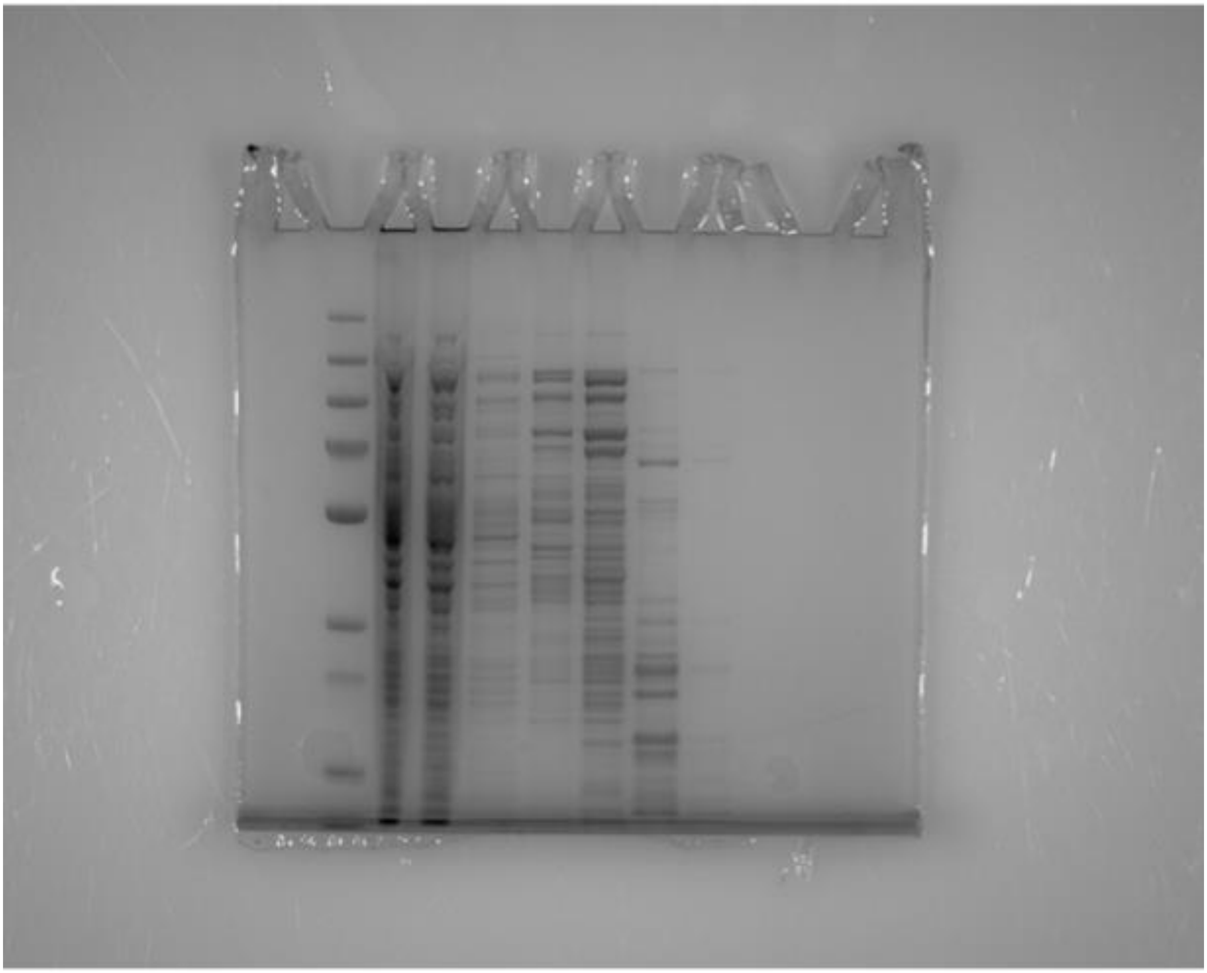

Supplementary Figure 8b – Full Western blot membrane

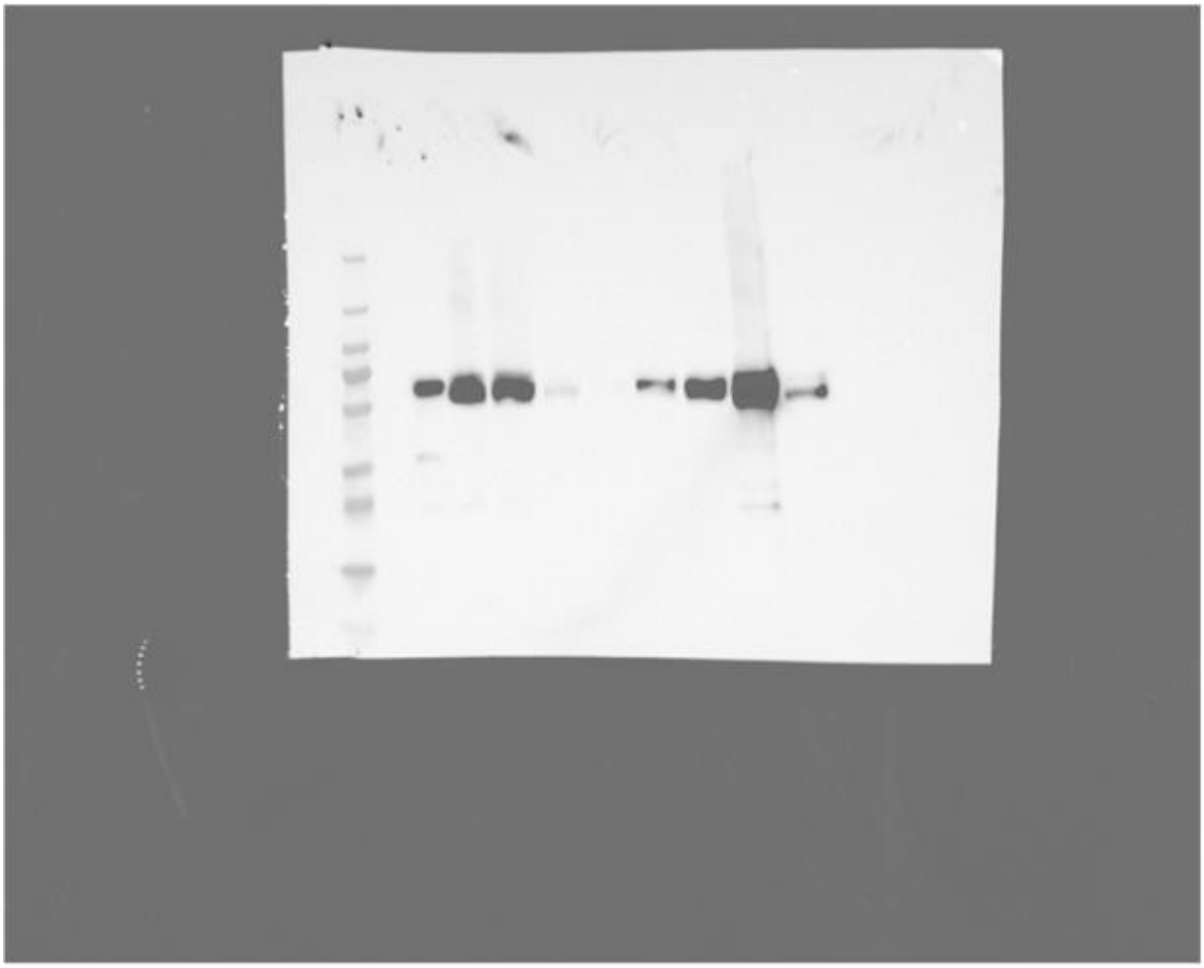
